# Transport efficiency of membrane-anchored kinesin-1 motors depends on motor density and diffusivity

**DOI:** 10.1101/064246

**Authors:** Rahul Grover, Janine Fischer, Friedrich W. Schwarz, Wilhelm J. Walter, Petra Schwille, Stefan Diez

## Abstract

In eukaryotic cells, membranous vesicles and organelles are transported by ensembles of motor proteins. These motors, such as kinesin-1, have been well characterized *in vitro* as single molecules or as ensembles rigidly attached to non-biological substrates. However, the collective transport by membrane-anchored motors, i.e. motors attached to a fluid lipid bilayer, is poorly understood. Here, we investigate the influence of motors anchorage to a lipid bilayer on the collective transport characteristics. We reconstituted ‘membrane-anchored’ gliding motility assays using truncated kinesin-1 motors with a streptavidin-binding-peptide tag that can attach to streptavidin-loaded, supported lipid bilayers. We found that the diffusing kinesin-1 motors propelled the microtubules in presence of ATP. Notably, we found the gliding velocity of the microtubules to be strongly dependent on the number of motors and their diffusivity in the lipid bilayer. The microtubule gliding velocity increased with increasing motor density and membrane viscosity, reaching up to the stepping velocity of single-motors. This finding is in contrast to conventional gliding motility assays where the density of surface-immobilized kinesin-1 motors does not influence the microtubule velocity over a wide range. We reason, that the transport efficiency of membrane-anchored motors is reduced because of their slippage in the lipid bilayer, an effect which we directly observed using singlemolecule fluorescence microscopy. Our results illustrate the importance of the motor-cargo coupling, which potentially provides cells with an additional means of regulating the efficiency of cargo transport.

## Significance

Molecular motors, such as kinesin-1, are essential molecules involved in active intracellular transport. Current mechanistic insights on transport by motors are mainly based on *in vitro* studies where the motors are bound to rigid substrates. However, when transporting membranous cargo under physiological conditions, multiple motors are often only loosely coupled via a lipid bilayer. In this study, we investigate how the motors’ transport efficiency is affected when bound to a lipid bilayer. In our reconstituted gliding motility assays, we show that membrane-anchored motors exhibit reduced transport efficiency due to slippage in the lipid bilayer. Notably, the efficiency increases at higher motor density and reduced membrane diffusivity, providing cells with an additional means of regulating the efficiency of cargo transport.

Intracellular transport of membrane-bound vesicles and organelles is a process fundamental to many cellular functions including cell morphogenesis, signaling and growth (1–4). Active cargo transport inside the eukaryotic cells is mediated by ensembles of motor proteins, such as kinesins and dynein, walking on microtubule tracks (5), and myosins walking on actin filaments (6). Gaining mechanistic insight into the functioning of these motors inside the complex environment of cells is challenging. Several studies have thus employed *in vitro* approaches to investigate the transport mediated by groups of same or different motors attached to cargos such as silica beads (7), quantum dots (8), glass coverslips (9, 10) or DNA scaffolds (11, 12). Although, these approaches provide us with knowledge about the collective dynamics of multi-motor transport, a key anomaly in these *in vitro* systems is the use of non-physiological rigid cargo. Vesicular cargo transport by molecular motors entails their attachment to fluid lipid bilayer either directly or via different adaptor molecules. The anchoring of motors in a diffusive lipid environment induce loose inter-motor coupling and the motors can diffuse within the lipid bilayer, thereby increasing the flexibility of the system. The effect of membrane-motor coupling on the transport behavior of motors is poorly understood. In addition, not much is known about how the transport efficiency of the motors is affected by their density and diffusivity on the membranous cargo.

In recent past, a few *in vitro* studies have been performed with membranous cargo such as liposomes (13) or supported lipid bilayers (SLBs) (14) to investigate the transport characteristics of actin filaments based motor proteins such as myosin Va and myosin 1c. Moreover, a recent study from our group reported the long-range transport of giant vesicles (1-4 μm) driven by kinesin-1 motors as a proof of concept that model membrane systems can by utilized to study microtubule-based motors such as kinesin-1 (15). However, due to the spherical geometry of GUVs or liposomes, it is difficult to determine unequivocally the number of motors transporting the cargo. Therefore, investigating the cooperative effects in cargo transport driven by membrane-anchored motors in such membrane-model systems is difficult. To circumvent this problem, we reconstituted the transport driven by multiple-motors anchored to lipid bilayer in an inverse geometry. We anchored recombinant truncated kinesin-1 motors with streptavidin binding peptide tag (SBP) to a flat biotinylated supported lipid bilayer (SLB) via streptavidin. These membrane-anchored motors were found to propel the microtubules in the presence of ATP. We examined the transport efficiency of such motors (ratio of observed microtubule gliding velocity and maximal microtubule gliding velocity) at different motor density and diffusivity. The flat geometry of the lipid bilayer allowed us to directly determine the density and diffusivity of GFP-tagged kinesin-1 using total internal reflection fluorescence (TIRF) microscopy. We observed that the transport efficiency of membrane-anchored kinesin-1 motors is reduced due to motor slippage in the membrane. However, the transport efficiency is increased at higher density and reduced diffusivity of motors in the membrane.

## Results

**Diffusivity of kinesin-1 motors anchored to SLB is determined by the diffusivity of lipids in SLB.** SLBs are one of the most suitable model membrane systems to quantify protein-lipid interactions (14, 16, 17). They can be subjected to single molecule imaging techniques such as TIRF microscopy, due to their flatness. This is not possible with other membrane-model systems such as giant unilamellar vesicles or liposomes due to their spherical geometry. Using TIRF microscopy we could quantify the density of motors as well as the motor diffusivity on SLBs with high accuracy. SLBs with 1,2-dioleoyl-sn-glycero-3-phosphocholine (DOPC) and 1,2-distearoyl-sn-glycero-3-phosphoethanolamine-N-[biotinyl(polyethyleneglycol)-2000] (DSPE-PEG-2000-Biotin) in molar ratio 99:1 were prepared by fusion of small unilamellar vesicles on hydrophilic glass coverslip. This lipid mixture was doped with 0.05 % 1,2-dioleoyl-sn-glycero-3-phosphoethanolamine-Atto647n (DOPE-ATTO647n) as a fluorescent marker. Homogenous SLBs were obtained (see Methods) as indicated by the uniform spreading of lipid marker, visible under the fluorescence microscope (Fig. 1A). Fluorescence recovery after photobleaching (FRAP) experiments were performed to determine the fluidity of the lipids in SLBs (see Methods). The diffusion coefficient of lipids in SLB was obtained to be *D*_Lipid_ = 3.0 ± 0.3 μm^2^/s (*N* = 16 SLBs, four independent experiments, Fig. 1B) by fitting the fluorescence recovery curves, according to the methodology described in Ref. (18).

**Figure 1.**
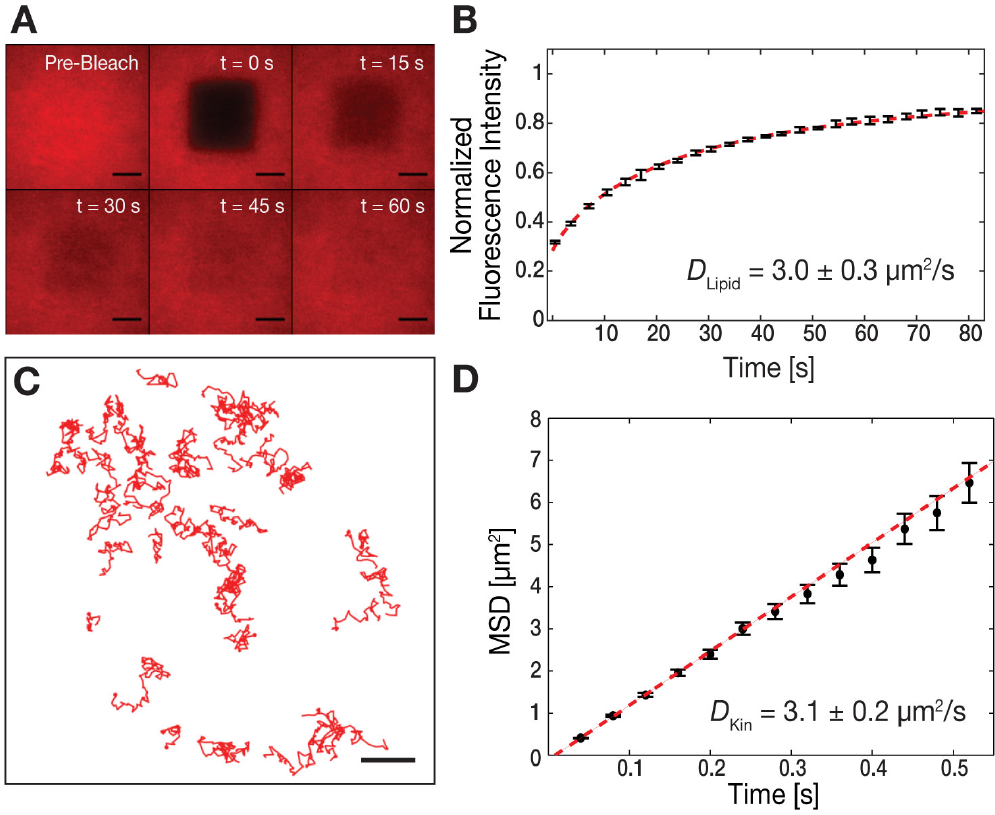
Membrane-anchored kinesin-1 motors diffuse with similar diffusivity as the lipids. (A) Time-lapse series of fluorescence recovery after photobleaching (FRAP) images for a SLB with a molar composition of DOPC:DSPE-PEG-2000-Biotin:DOPE-Atto647n = 99:1:0.05 (Scale bar, 10 μm). (B) Representative curve for the normalized fluorescence intensity after photobleaching vs. time (black, *N* = 4 SLBs, mean ± SD). The diffusivity of the lipid bilayer was determined to be Dypid = 3.0 ± 0.3 μm^2^/s (mean ± SD, *N* = 4 independent experiments, dashed red line, fit according to Ref. (18)). (C) Single-molecule trajectories of freely diffusing rKin430-SBP-GFP anchored on a biotinylated SLB via streptavidin (Scale bar, 10 μm). (D) MSD data (black, mean ± SD) of diffusing rKin430-SBP-GFP molecules. The diffusion coefficient was determined to be *D*_kid_ = 3.1 ± 0.2 μm^2^/s (mean ± 95% confidence interval, *N* = 40 tracked molecules, red line, linear fit to the first eight points).

We expressed and purified rat kinesin-1 heavy chain isoform KIF5C, truncated to 430 a.a (19), with SBP tag at the tail (rKin430-SBP) (20), as well as with SBP and GFP-tag at the tail (rKin430-SBP-GFP). rKin430-SBP was anchored to biotinylated SLB via streptavidin. Thereby, biotinylated SLBs were incubated with a saturating concentration of streptavidin (0.5 μM), 100 folds higher than the number of biotinylated lipids. Thus, it was ensured that the surface density of motors on the streptavidin loaded biotinylated SLBs is regulated by the bulk concentration of motors applied to the reaction chamber and not by the number of available binding sites. To measure the diffusivity of motors anchored to SLB we added GFP-labeled rKin430-SBP at low concentration (<10 nM) along with rKin430-SBP (0.25 μM) to the streptavidin-coated biotinylated SLBs. The diffusing single molecules of rKin430-SBP-GFP were excited with a 488-nm laser and imaged using TIRF microscopy. Single-particles were tracked using the Fluorescence Image Evaluation Software for Tracking and Analysis (FIESTA) (21) to obtain the single molecule trajectories (Fig. 1C). The ensemble average diffusion coefficient of rKin430-SBP-GFP was obtained to be *D*_Kin_ = 3.1 ± 0.2 μm^2^/s (mean ± 95% confidence interval, *N* = 40 molecules) as characterized by linear fitting to the mean square displacement (MSD) as a function of time (Fig. 1D, see Methods). The diffusion coefficient of rKin430-SBP obtained from singleparticle tracking is consistent with the fluidity of lipids in SLBs obtained from FRAP analysis. Thus, our results validate that the diffusivity of motors attached to a SLB is governed by the fluidity of lipid anchors in the SLB. Furthermore, the single-particle tracking studies confirm that the rKin430-SBP is freely diffusing upon anchoring to SLBs.

**Microtubule gliding velocity increased with increasing surface density of membrane-anchored motors.** It has been previously reported that the microtubule gliding velocity propelled by surface-immobilized kinesin-1, is independent of the motor density over a wide range (22, 23). But are the transport characteristics of loosely coupled membrane-anchored motors different from the surface-immobilized motors? To address this question, we reconstituted microtubule gliding driven by kinesin-1 anchored to a SLB. Experimental set-up of the membrane-anchored gliding assay is shown in (Fig. 2A). Rhodamine-labeled, double-stabilized microtubules (see Methods) were applied to the membrane-anchored rKin430-SBP, in a buffer solution with 1 mM ATP. The motors anchored to the SLB propelled the microtubules, confirming that the motors were functional upon binding to biotinylated SLBs. We systematically varied the surface density of motors on the SLBs by incubating streptavidin-bound biotinylated SLBs with different concentration of rKin430-SBP ranging from (0.07 μM - 0.63 μM). We observed smooth microtubule gliding at high motor densities, however, microtubule translocation got slower and wigglier at lower motor densities (Movie S1).

**Figure 2.**
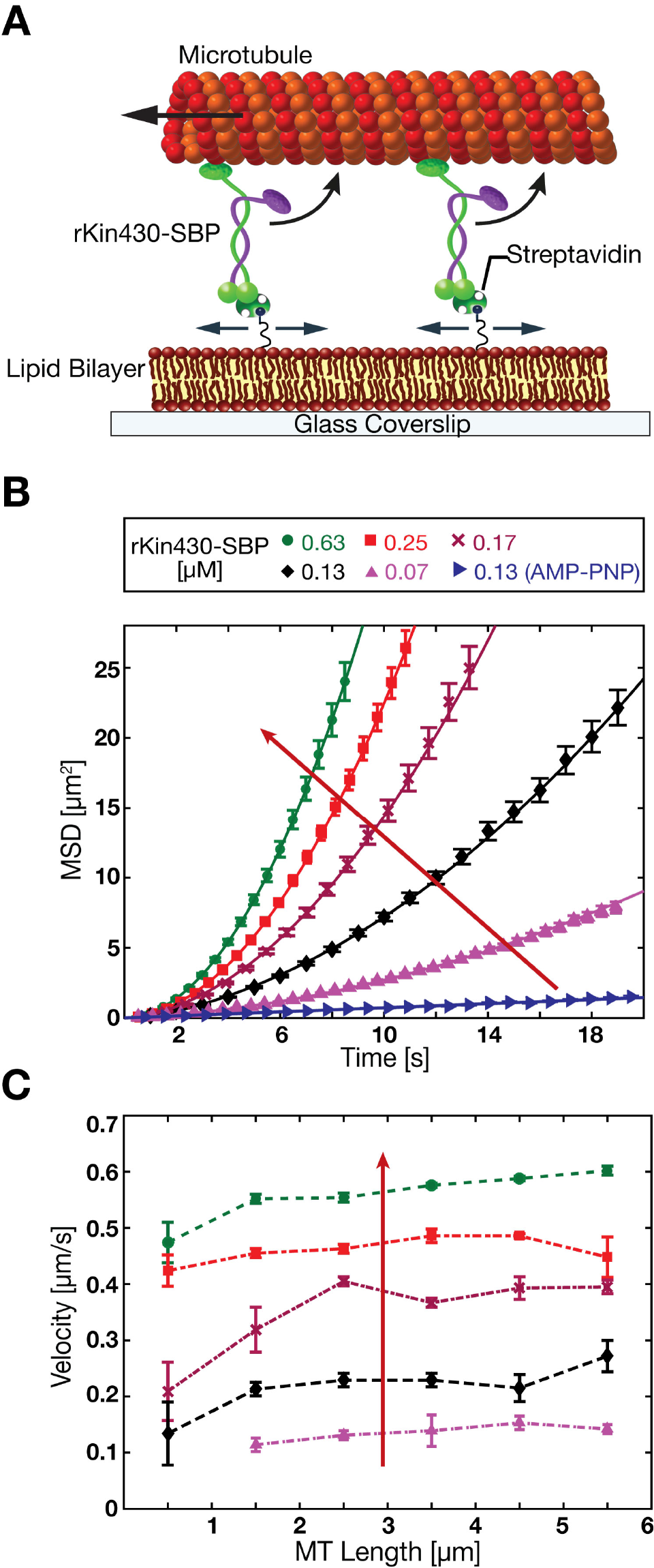
In-vitro reconstitution of a membrane-anchored gliding motility assay. (A) Schematic drawing (not drawn to scale) of the experimental setup: truncated rat kinesin-1 with streptavidin-binding-tag (rKin430-SBP) is attached, via streptavidin, to a biotinylated SLB. The motors diffusively-anchored on the SLB propel the microtubules. (B) Representative ensemble MSD data for the center positions of the microtubules (mean ± SEM, *N* ≥ 40 microtubules) at different motor concentrations in 1 mM ATP and 1mM AMP-PNP (only for 0.13 μM rKin430-SBP). The red arrow indicates increasing motor concentration. To calculate the linear translocation components (i.e. the microtubule velocities v) the data was fit by Eq. 1. (C) Ensemble-averaged microtubule gliding velocities for different microtubule lengths, binned into 1 μm intervals, at different motor concentrations (mean ± 95% confidence interval, *N* ≥ 15 microtubules for each data point).

To calculate the microtubule gliding velocity, translocation of microtubules was divided into two components (i) translational component due to active transport by the motors and (ii) diffusional component due to attachment to a diffusive lipid bilayer via motors. Translational and diffusive component were determined by fitting the MSD of the microtubule center over time with the following equation as described in (24)

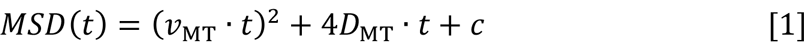

here *v*_MT_ is the translational velocity of a microtubule, *D*_MT_ is the diffusional component and *c* is the offset accounting for the localization uncertainty and the dynamic error due to finite camera acquisition time (25). We validated that the microtubules driven by membrane-anchored motors have both translational and diffusional component by performing motility assay in presence of 1 mM adenylyl imidodiphosphate tetralithium (AMP-PNP), a non-hydrolysable analog of ATP. This ensured that the motors were bound to the microtubules in a rigor state, thus the microtubules only have diffusional component. In fact, this is what we observed (Movie S1), where the MSD of the microtubules in AMP-PNP increased only linearly with time (Fig. 2B).

The MSD data of the gliding microtubules fitted well to the Eq. 1 for all motor concentrations (Fig. 2B). The slope of the MSD curves vs. time increased with increasing motor concentration. To resolve whether the microtubule gliding velocity was dependent on the number of motors or the motor density, we binned the microtubules over 1 μm length intervals and determined the average microtubule velocities. We observed that the gliding velocities were largely independent of the microtubule lengths for all motor concentrations (Fig. 2C). Thus, the transport efficiency of the membrane-anchored motors propelling a microtubule is set by the number of motors per unit length of a microtubule and not by the absolute number of motors.

**Membrane anchored motors slip backward in lipid bilayer while propelling a microtubule forward**. One of the striking observations in our membrane-anchored gliding motility assays was that the microtubules upon collision did not cross each other (Fig. 3A upper panel, Movie S2). Taxol-stabilized GTP-grown microtubules always aligned with passing microtubules upon collision. Stiffer, double-stabilized microtubules aligned with passing microtubules when colliding at shallow angles, but temporarily stalled until the passing microtubules glided away when colliding at steep angles. This behavior is in contrast to gliding motility assays with surface-immobilized kinesin-1, where the microtubules cross over each other upon collision, without any noticeable hindrance (Fig. 3A lower panel, Movie S2, and Ref. (26)). We therefore hypothesize that the force output of membrane-anchored motors is lowered due to their membrane-anchored tails slipping in the lipid bilayer.

**Figure 3.**
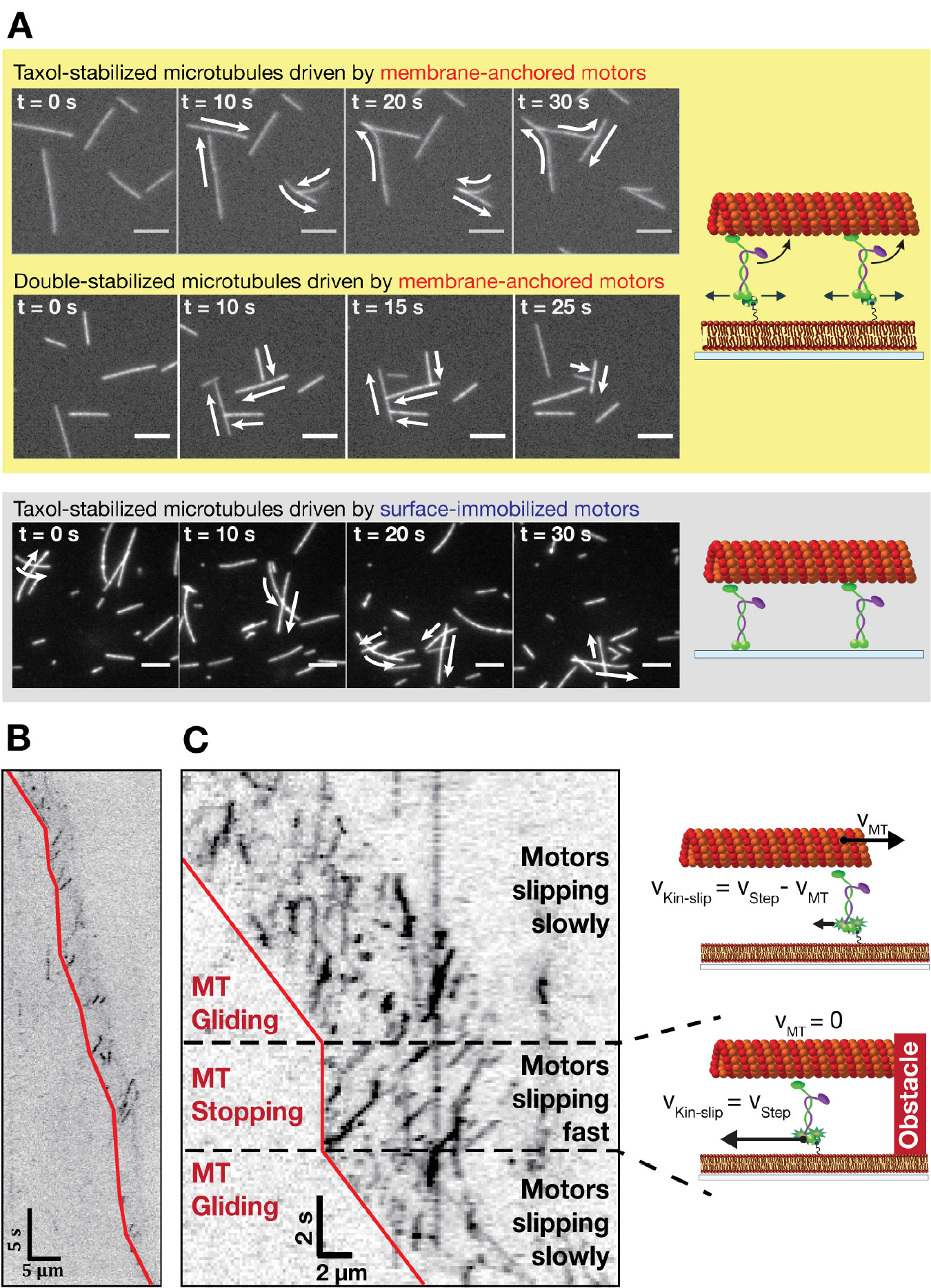
Membrane-anchored kinesin-1 motors slip in the lipid bilayer, while propelling a microtubule. (A) Time-lapse images for microtubules driven by membrane-anchored motors (upper panel) and surface-immobilized motors (lower panel) with schematic experimental setups on the right (Scale bar, 5 μm). The arrows indicate the transport directions of the gliding microtubules. Microtubules propelled by membrane-anchored motors do not cross each other in contrast to microtubules driven by surface-immobilized motors. (B-C) Representative kymographs (inverted contrast) showing the movement of individual rKin430-SBP-GFP motors (dark signals) while propelling microtubules at low motor density (B) and high motor density (C). The red lines mark the trailing ends of the microtubules as guides to the eye. In (C) a gliding microtubule collides with another passing microtubule and is temporarily stalled (also shown in the schematics on the right) until the other microtubule glides away.

To directly image whether the motors slip in the lipid bilayer while propelling a microtubule, we performed membrane-anchored gliding assays with rKin430- SBP spiked with low concentration (<25 nM) of rKin430-SBP-GFP. Spiking experiments enabled us to resolve single motors propelling a microtubule for a wide range of motor concentrations. We observed that at high motor density (when the microtubules moved fast) the membrane-anchored motors propelling it slipped backward slowly. However, at low motor density (when the microtubules moved slowly) the motors slipped backward at higher velocities (Fig. 3B). The relation between slipping velocity of motors in the lipid bilayer and microtubule gliding velocity in our experiments can be clearly seen when a fast moving microtubule encounters an obstacle (another passing microtubule) briefly, thus causing the microtubule to stall. When a moving microtubule is stalled, the motors underneath the microtubule started slipping backward at high speed, as seen in the kymograph (Fig. 3C). These results suggest that the motor slipping velocity and the microtubule gliding velocity are adding up to the stepping velocity of a single rKin430-SBP motor.

**Frictional forces on the membrane-anchored motors and on microtubule determine the transport efficiency.** To quantitatively describe our observation of increasing transport efficiency with increasing membrane-anchored motor density, we developed a theoretical description of membrane-anchored gliding motility. The nanoscopic set-up with the physical parameters used to develop the mathematical model is illustrated in (Fig. 4A). We resolved the dynamics of membrane-anchored gliding motility by considering the velocities and frictional forces of membrane-anchored motors and microtubules. (1) Velocities: Membrane-anchored motors step on a microtubule with a velocity 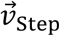 thus propelling a microtubule with a velocity 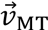 relative to the substrate, in a direction opposite to its stepping direction. As observed in the experiments, the motors drag their anchors in the lipid bilayer, under the microtubule, in the stepping direction with a velocity 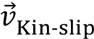 relative to the substrate, as the motor is not rigidly bound but anchored to a fluid bilayer. Thus, the relationship between the different velocities can be formulated as:

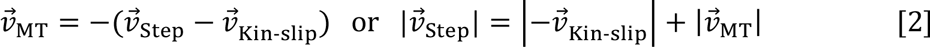

**Figure 4.**
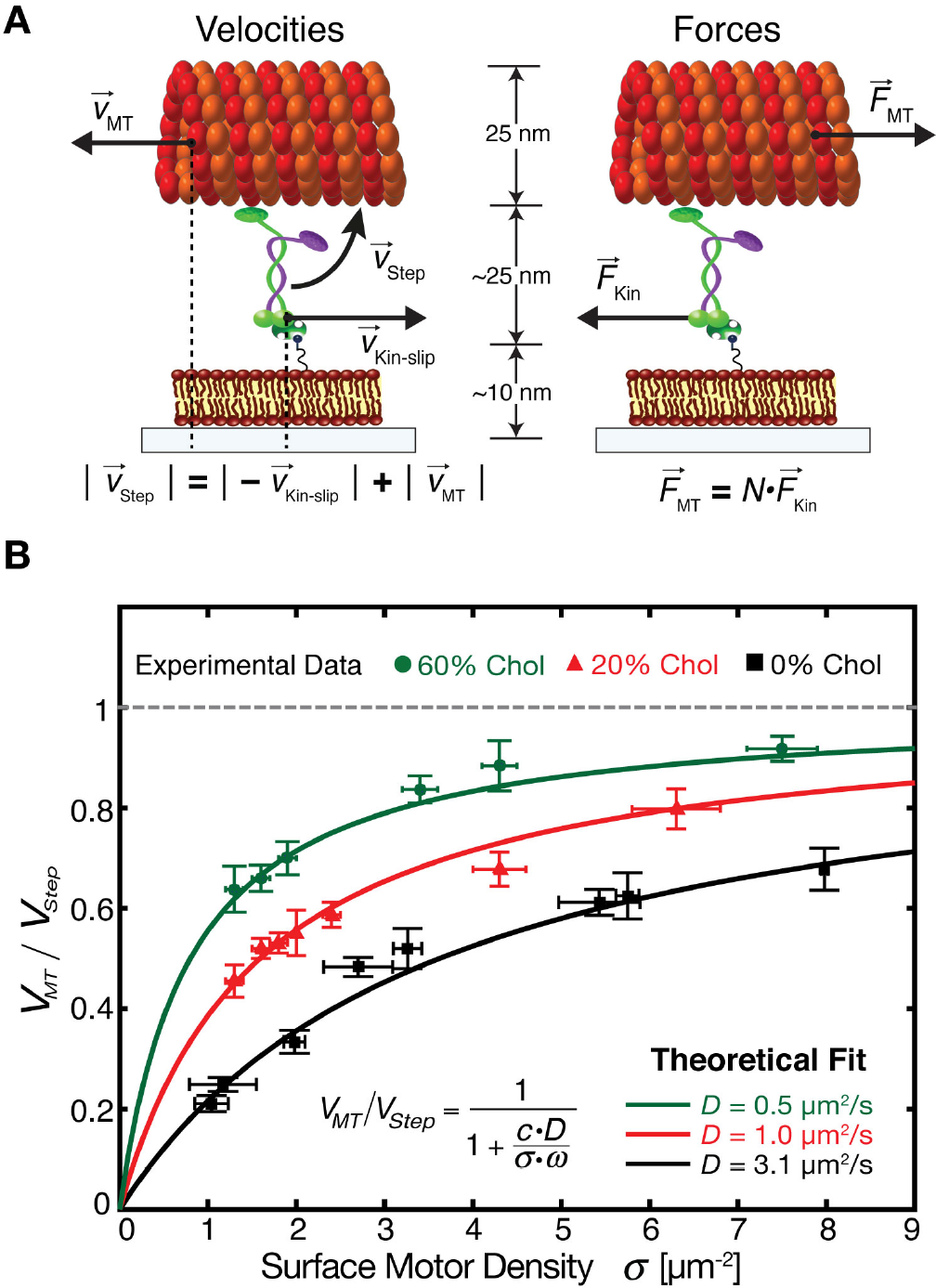
Theoretical description of the membrane-anchored gliding motility fits well to the experimental observations. (A) Nanoscopic view of the experimental set-up illustrating the velocities and frictional forces of microtubules and motors, used to derive the mathematical model. The depicted microtubules glide to the left with respect to the substrate at velocity vmt. The motor steps on the microtubule with velocity *v*_step_ and thereby moves its anchor in the SLB to the right at velocity *v*_Kin-slip_ with respect to the substrate. The frictional forces act on the microtubules and motors in the directions opposite to their motion. (B) Averaged microtubule transport efficiency (mean ± 95% confidence interval, *N* ≥ 40 microtubules for each data point) for various kinesin-1 surface densities (mean ± SD, *N* = 5 regions of interest) and different SLB compositions (0% CH, 20% CH and 60% CH). The data were fitted (solid lines) to the mathematical model (Eq. 13), with one free parameter *ω* (reach of a diffusing kinesin-1 motor to bind to a microtubule, *c* = 0.72 μm^−3^·s, *R*^2^ ≥ 0.95).

The stepping velocity of rKin430-SBP-GFP was determined by performing standard stepping motility assays, where the movement of motors on surface-immobilized microtubules in the assay buffer was recorded and analyzed (Methods, Fig. S1) with 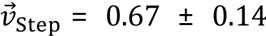 μm/s (mean ± SD, *N* = 545 molecules). (2) Forces: Under steady state, the net force acting on the system is zero, i.e. the forces on the motors balance the forces on the microtubule. The frictional force acting on a microtubule 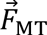 can be estimated by the hydrodynamic drag, due to its motility in the aqueous solution. The frictional force on a kinesin motor 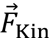 can be estimated by the drag in the fluid bilayer. At any instance there would be several motors *N* interacting with a microtubule. These *N* motors stepping, with the same velocity 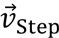, in an uncorrelated manner on a microtubule would experience equal drag force. The force balance equation then yields:

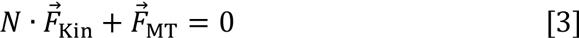

The frictional force on a microtubule can be estimated by considering it as a long rigid cylinder with its length much larger than its radius *L*_mt_ ≫ *T*_mt_. The drag coefficient for cylindrical objects moving parallel to the surface is given by (22)

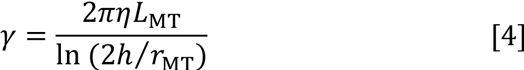

The magnitude of the drag force on the microtubule can be determined by the Stokes’ law:

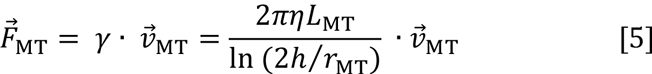

A microtubule would experience maximum drag force when its moving at highest velocity which is 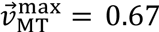 μm/s calculated from equation (2) considering 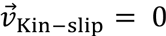. Thus for η = 10^−3^ Pa·s (water), *L*_MT_ = 10 μm, *h* = 50 nm and *r*_MT_ = 12.5 nm, the maximum force on a microtubule would be 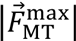 = 20 fN. The magnitude of maximum drag force on a microtubule in aqueous environment is 200-folds less than the stall force of a single kinesin-1 motor 5-7 pN (27, 28). Thus, even a single surface-immobilized kinesin-1 motor can propel a microtubule at maximum velocity in aqueous environment. Therefore, the microtubule gliding velocity is independent of motor density for surface-immobilized kinesin-1 over a wide range.

Frictional force on a membrane-anchored motor can be estimated by using the Einstein-Smoluchowski-relation where the drag coefficient is related to the diffusivity *D*_Kin_ of a molecule by

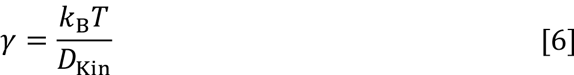

The magnitude of the force then can be determined by

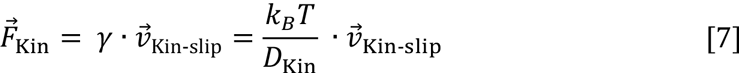

Single membrane-anchored kinesins would experience a maximum drag force when they are slipping under a stationary microtubule at its maximum stepping velocity, which is 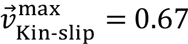 μm/s calculated from equation (2) considering 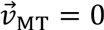. When interacting with a microtubule, the mobility of a motor is reduced to only one dimension since the rKin430-SBP head walk on a single protofilament of a microtubule. Thereby, the diffusion coefficient of microtubule-interacting motors in the lipid bilayer will be reduced to half. For *k*_B_ = 1.38 × 10^−23^ *J/K, T* = 295 *K* and *D*_Kin-1D_ = *D*_Kin_/2 = 1.56 μm^2^/s, the maximum force on membrane-anchored rKin430-SBP propelling a microtubule would be 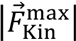 = 1.7 fN. This value is 250 folds smaller than the stall force of a single kinesin-1 motor. Thus, kinesin-1 motors under the low load condition of our set-up would step on a microtubule at their maximum stepping velocity. By substituting the expression of forces for microtubule and kinesin motor in the equation (3) we obtain

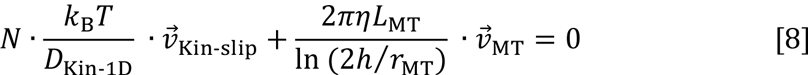

which can be simplified to

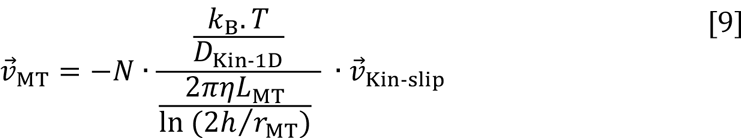

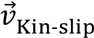 is substituted in the above equation as 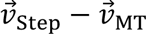 from equation (2), to obtain the relation

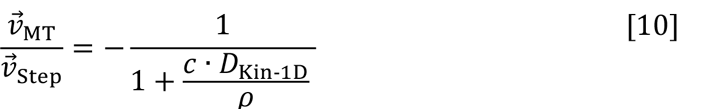

where 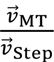 is the transport efficiency, ρ = *N*/*L*_MT_ is the number of motors interacting with a microtubule per unit length i.e. the linear motor density, and *c* is a constant which depends on the physical parameters given by

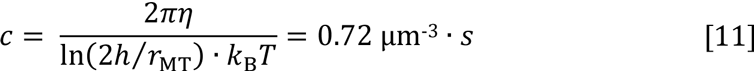

Hereby, we have derived a simple model that describes the dependence of the transport efficiency of membrane-anchored motors on their linear density and their diffusivity. The model predicts that the transport efficiency of membrane-anchored motors increases with increasing motor density but is independent of the microtubule length which is what we observed in our membrane-anchored gliding assays (Fig. 2B-C). For motors immobilized on the glass substrate, the apparent *D*_Kin-1D_ is zero. Substituting into Eq. 10, we obtain that the microtubule gliding velocity is as high as the single motor stepping velocity and is independent of the motor density. This result has been reported previously for kinesin-1 (23, 29) and what we observe for rKin430-SBP over a wide range of motor density. However, for the membrane-anchored motors the model predicts higher transport efficiency at high motor density and/or at low diffusivity of the motors’ lipid anchors.

**Theoretical model describes the experimental findings well.** We experimentally tested the predictions of our theoretical model, i.e. the dependence of transport efficiency of membrane-anchored motors on i) motor density on SLBs and ii) diffusivity of motors lipid-anchor in SLB. To measure the surface density of motors on the SLBs, we incubated the streptavidin-coated biotinylated SLBs with rKin430-SBP along with a low concentration of GFP-labeled rKin430-SBP (~20 nM), in a fixed molar ratio of 150:1. The samples were excited at 488 nm and the images of GFP-labeled single motors diffusing on the SLBs were recorded, using TIRF microscopy. To avoid errors in counting due to photobleaching, we only considered the first five frames of the movie stream to determine the number of motors anchored to a SLB (see Methods). The motor surface density was determined by averaging the total number of diffusing particles, per unit area of the field of view. Using this methodology we could directly measure the surface density of motors. Subsequently, membrane-anchored gliding motility assays were performed on the same samples to obtain the microtubule gliding velocity, for a measured motor surface density. To compare our experimental data with the theoretical model, we assumed a linear relationship between the surface motor density *σ* and the linear motor density *ρ*, given by (30)

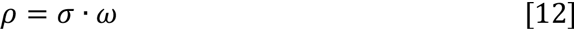

where *ω* is the interaction reach of the motor to bind to a microtubule filament. Thus the equation [10] can be written as

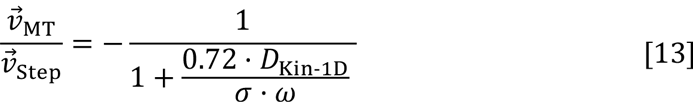

The experiments were performed over a wide range of surface motor density to quantify the dependence of transport efficiency on motor density (Fig. 4B).

Our theoretical model predicts that at a fixed motor density, the transport efficiency of membrane-anchored motors increases with increasing viscosity of the lipid anchors. To test this prediction, we varied the diffusivity of the SLBs to which the motors were anchored, by adding cholesterol. It has been previously shown that the addition of cholesterol increases the packing of lipids with unsaturated acyl chains such as DOPC and thus reduces the diffusivity of SLBs (31). We added cholesterol to the lipid mixture in two different molar ratios (20% and 60% of the total lipid concentration) with the composition D0PC:Cholesterol:DSPE-PEG-2000-Biotin 79:20:1 (20% CH) and DOPC: Cholesterol:DSPE-PEG-2000-Biotin 39:60:1 (60% CH), along with a trace amount of fluorescent lipid-marker DOPE-ATTO647n to check the fluidity and homogeneity of SLBs. We performed FRAP experiments and found that the diffusivity of the SLBs reduced after addition of cholesterol, with 60% CH being less diffusive than 20% CH (Movie S3). Subsequently, we measured the diffusivity of membrane-anchored motors (rKin430-SBP along with a low concentration of rKin430-SBP-GFP) in the CH SLBs (Methods) and obtained 1.0 ± 0.2 μm^2^/s; (mean ± 95% confidence interval; *N* = 48 molecules) and 0.5 ± 0.1 μm^2^/s (mean ± 95% confidence interval, *N* = 58 molecules) for 20% and 60% CH SLBs, respectively. Thus, with single molecule tracking of rKin430-SBP-GFP we confirmed that the addition of cholesterol in our experimental set-up reduced the diffusivity of the membrane-anchored motors.

Subsequently, membrane-anchored gliding motility assays at different motor densities were performed on the CH SLBs. We observed that, on 20% CH SLBs microtubules did not cross each other upon collision, whereas on 60% CH SLBs they did (Movie S4). Thus, the motors anchored to 60% CH SLBs were less slippery than the motors anchored to 20% CH SLBs and therefore produced enough force output to propel microtubules over each other. The transport efficiency of motors anchored to CH SLBs also increased with increasing motor density, similar to our findings on 0% CH SLBs (Fig. 4B). Furthermore, for a given motor density, the transport efficiency was highest for 60% CH SLBs followed by 20% CH and least for 0% CH SLBs. We fitted our experimental data with the theoretical model (Eq. 13) using fitting routines in MATLAB (Methods). Our theoretical model with only one free parameter (namely *ω*) fitted well to the experimental data for different motor densities as well as different diffusivities (Fig. 4B). From the fits we obtained the values of *ω* to be 0.31 ± 0.07 μm, 0.24 ± 01 μm, and 0.22 ± 0.02 μm, (mean ± 95% confidence interval) for 0% CH, 20% CH and 60% CH respectively. Previous studies have estimated the value of *ω* for kinesin-1 motors immobilized on a solid substrate to be around 0.02 μm (30). However, for our experimental set-up we expect a higher reach as the kinesin-1 motors are diffusing freely on a lipid bilayer. For example a membrane-anchored kinesin-1 diffusing at 3.1 μm^2^/s can explore a circle of radius 0.45 μm in 50 ms. Thus, the obtained value of ω is reasonable for our membrane-anchored gliding motility assays. Our experimental data is consistent with the predictions from our theoretical model, describing the increase in transport efficiency of membrane-anchored motors with increasing motor density and increasing viscosity.

## Discussion

In this study, we established membrane-anchored gliding motility assays, to investigate the transport characteristics of kinesin-1 motors that are loosely coupled via a lipid bilayer. For kinesin-1 tightly coupled to a substrate it has been previously reported that the microtubule gliding velocity is either independent of the motor density or reduced at higher motor density (23, 32). Our results show that the gliding velocity of microtubules driven by membrane-anchored kinesin-1 is reduced due to slipping of the motor anchors in the lipid bilayer. However, the gliding velocity increases with increasing motor density or increasing viscosity.

*In vivo*, cargo transport velocities determined by tracking vesicles or organelles in various cell types exhibit significant spreads and the velocity histograms often contain multiple peaks (33–36). In the complex environment of a cell, there could be many factors that influence cargo transport, such as motor activity, membrane-motor binding partners, the movement of cytoskeletal filaments (37) etc. to produce faster or slower motility. One way in which the transport can be regulated inside cells is by varying the lipid composition of membranous cargos. For example, if the motors are segregated into lipid-microdomains, both the frictional forces on the motor anchors as well as the motor density would increase. This results in less slippage of the motors and thus an increased transport efficiency. Along these lines, a recent study has reported that cytoplasmic dynein clusters into lipid-microdomains on phagosomes to drive rapid transport towards lysosomes (38). The importance of motor density and membrane rigidity was also highlighted previously when tubular transport intermediates between organelles were reconstituted by extracting nano-vesicular tubes from GUVs with kinesin-1 motors (39, 40). There it was shown that a minimum motor density was required to pull the membrane tubes from GUVs depending on the lipid composition.

To the best of our knowledge, this is the first reconstitution of microtubule gliding motility on membrane-anchored motor proteins. We believe, our assay provides a useful tool to study the regulation of cargo transport by a variety of molecular motors such as kinesin-3 (KIF16B, KIF1A), which can directly bind to a membranous cargo. Furthermore, these assays can be used to gain mechanistic insight into the recruitment of various motors to their specific cargo and their transport characteristics. So far, our experiments were performed on flat membranes and do not yet account for varying cargo geometries such as spherical vesicles (as in secretory granules) and small tubules (as in the Endoplasmic reticulum). Most likely, this geometry additionally influences the transport efficiency and will be a subject of our further studies.

## Methods

**Supported lipid bilayer (SLB) preparation.** Glass coverslips were prepared by 15 min sonication in 5% Mucasol solution, extensive rinsing with nanopure water, 15 min sonication in 100% ethanol, and again extensive rinsing with nanopure water. Coverslips were dried with pressured nitrogen and plasma cleaned (Diener electronic) for 10 min. Multi-lamellar vesicles (MLVs) of the desired lipid mixture were prepared by taking 7.5 μg of total lipids in a glass vial (Sigma) in the required molar ratio and evaporating the solvent (chloroform) under a constant stream of nitrogen. Any residual solvent was removed by keeping the glass vial under vacuum overnight. The lipids were then rehydrated in H20S75 buffer (20 mM HEPES, 75 mM NaCl, pH 7.2) to a concentration of 0.2 mg/ml and sonicated for 20 min to form small unilamellar vesicles (SUVs). The SUV dispersion was added to the experimental chambers, formed by attaching a cut 200 μl Eppendorf tube using UV adhesive (NOA 83, Norland products) to a plasma cleaned glass coverslip. CaCh was added to a final concentration of 3 mM to induce fusion of SUVs and formation of an SLB. After 45 min of incubation at room temperature, the sample was washed with 1 ml of H20S75 buffer in steps of 50 μl to remove unfused vesicles. SLBs were prepared with 1,2-dioleoyl-sn-glycero-3-phosphocholine (DOPC, Avanti) and 1,2-distearoyl-sn-glycero-3- phosphoethanolamine-N-[biotinyl(polyethylene-glycol)-2000] (DSPE-PEG-2000- Biotin, Avanti) in molar ratio 99:1 with 0.05% 1,2-dioleoyl-sn-glycero-3-phosphoethanolamine-Atto647n (DOPE-ATTO647n, Atto-Tec fluorescent labels) as a fluorescent lipid marker. The composition of SLBs with cholesterol was D0PC:Cholesterol(Avanti):DSPE-PEG-2000-Biotin = 79:20:1 (20% CH) and D0PC:Cholesterol:DSPE-PEG-2000-Biotin = 39:60:1 (60% CH).

**rKin430-SBP purification and attachment to SLBs.** A codon optimized DNA sequence of kinesin-1 containing the N-terminal 430 amino acids of the *Rattus norvegicus* kinesin-1 isoform kif5c, with the tags 8xHis, mCherry and SBP, was purchased from Invitrogen (GeneArt, Invtirogen). Two restriction sites, PacI and AscI, were introduced in pET24d vector (#69752-3, Addgene) and the rKin430- mCherry-SBP sequence was inserted in the vector after digesting it with PacI and AscI restriction enzymes (New England Bioloabs). The mCherry sequence was cut out using the restriction enzyme NgoMIV and the cut plasmid was ligated to obtain the rKin430-SBP plasmid. The rKin430-SBP-GFP DNA sequence was prepared by inserting a multifunctionalGFP (mfGFP) tag (20) having 8xHis, SBP, and c-Myc tag, in tandem in a loop of the GFP sequence. The sequence was inserted into rKin430 plasmid (19) in pET17 vector (# 69663-3, Addgene). Briefly, PCR with primers GGTACCGTGAGCAAGGGCGAGGAGCTGTTC and CAATTGTTACTTGTACAGCTCGTCCATGCCGAGAGTG was used to amplify the mfGFP sequence and restriction sites KpnI and MfeI were added at the 5’ and 3’ end of the complementary sequence, respectively. The mfGFP sequence was then inserted into rKin430 plasmid in pET17 vector using restriction enzymes KpnI and MfeI (New England Bioloabs). Both constructs were expressed in BL21 (DE3) *E. coli* cells. Expression with isopropyl (β-D-1-thiogalactopyranoside (IPTG) and affinity purification with the HisTrap Column (GE Health Care) was carried out as described previously (19). rKin430-SBP and rKin430-SBP-GFP concentrations were quantified by comparison with the known GFP standards on a Coomassie-stained 4-12% Bis-Tris NuPAGE gel (Thermo Fisher Scientific).

rKin430-SBP and rKin430-SBP-GFP were attached to SLBs, by first incubating the biotinylated SLBs with 0.5 μM streptavidin (Sigma) in 100 μl total volume for 10 min followed by washing with 1 ml of H20S75 buffer to remove the unbound streptavidin. Before adding the motors to the streptavidin-loaded SLBs, the experimental chambers were equilibrated by exchanging H20S75 buffer with motor buffer (20 mM HEPES, 75 mM NaCl, 1 mM ATP, 1 mM MgCh, 1 mM DTT and 20 mM D-glucose). 50 μl of the buffer in the experimental chamber were then replaced with 50 μl of motor solution, consisting of the required amount of rKin430-SBP or rKin430-SBP-GFP in motor buffer. After incubating the motors on the streptavidin-loaded SLBs for 6 min, unbound motors were washed with 200 μl of assay buffer (20 mM HEPES, 75 mM NaCl, 1 mM ATP, 1 mM MgCh, 1 mM DTT, 40 mM D-glucose, 0.02 mg/ml glucose oxidase, 0.01 mg/ml catalase and 1 μM taxol).

**Stepping and gliding motility assays.** Experiments to obtain the stepping velocity of single rKin430-SBP-GFP motors were performed in flow channels made of silanized coverslips, as described in (41). Briefly, first a solution of β-tubulin antibodies (0.5% SAP.4G5, Thermo Fisher Scientific) diluted in H20S75 buffer was flushed into a flow channel and incubated for 5 min, followed by a washing step with H20S75 buffer. The flow channel was then incubated with 1% Pluronic F127 in H20S75 for 45 minutes to block the surface from unspecific binding of proteins. Subsequently, the flow channel was washed with 80 μl of H20S75 supplemented with 10 μM taxol (H20S75T). Fluorescent, taxol-stabilized rhodamine microtubules (used in the stepping motility assays as well as for the data in Fig. 3A and B) were prepared as described previously (41). Taxol-stabilized GMP-CPP microtubules (also referred to as ‘double-stabilized microtubules’, used throughout our work unless mentioned otherwise) were prepared by polymerizing 2.5 μM tubulin mix (1:3 rhodamine labeled tubulin: unlabeled tubulin) in BRB80 buffer (80 mM PIPES, 1 mM MgCh, 1 mM EGTA, pH 6.9) supplemented with 1.25 mM GMP-CPP and 1.25 mM MgCh. The polymerization was carried out at 37 °C for 2 hours in 80 μl BRB80, after which 120 μl of BRB80 with 15 μM taxol is added. Microtubules were always prepared freshly before each experiment and free tubulin was removed by ultracentrifugation in an Airfuge (Beckman Coulter) at 100,000 × g for 10 minutes. The pellet was re-suspended in 200 μl H20S75T buffer. A solution of microtubules was flushed in and allowed to attach to the antibodies on the coverslip for 5 minutes. Unbound microtubules were removed from the flow channel by washing with 40 μl of H20S75T. Finally, 20 μl of 100 pM rKin430-SBP-GFP in assay buffer was flushed into the flow channel.

**Image acquisition for single-molecule TIRF and FRAP experiments.** Images were obtained using a Nikon microscope (Nikon Eclipse Ti equipped with Perfect Focus System (PFS) and a FRAP module, PlanApo 100x oil immersion objective lens, NA 1.49) with an electron multiplying charge-couple device (EMCCD) camera (iXon ultra EMCCD, DU-897U, Andor) in conjunction with NIS-Elements (Nikon) software. Single-molecule imaging for stepping assays, and singleparticle tracking of rKin430-SBP-GFP on SLBs were performed using TIRF microscopy and a monolithic laser combiner (Agilent MLC 400) which has a dual output for FRAP and fluorescence imaging. SLBs were imaged with a Cy5 filter set (exc: 642/20. Dichroic LP 647, em: 700/75) and rKin430-SBP-GFP molecules were imaged with another filter set (exc: 475/35. Dichroic LP 491, em: 525/45). Images were acquired in continuous streaming mode with 100 ms exposure for stepping assays and 50 ms exposure for single-particle tracking, to localize the diffusing motors on SLBs. For estimating the motor density on SLBs, GFP-labeled motors were imaged for 150 frames (256 × 256 pixels) with 100 ms exposure time each. For photobleaching experiments 512 × 512 pixel images of the SLBs were captured at 0.1 s interval for 1 s using the 647-nm laser line, following which a 150 x 150 pixel region in the center of field of view was bleached using 647-nm laser at full power using the FRAP module for 4.2 s (5 scan iterations). Time-lapse images were then recorded at an interval of 0.5 s for 250 frames to monitor the recovered fluorescence in the bleached area. Images of rhodamine-labeled microtubules in membrane gliding motility assays were observed by epi-fluorescence where microtubules were excited with a metal arc lamp (Intensilight, Nikon) in conjunction with a rhodamine filter set (exc: 555/25. Dichroic LP 561, em: 609/54).

**Data analysis for FRAP, single-particle tracking and motility assays.** FRAP images were analyzed to estimate the diffusion coefficient of the lipids in a SLB using an algorithm described in Ref. (18). The MATLAB script was modified to correct for the fixed pattern noise arising due to non-uniform illumination and TIRF imaging. To correct for the fixed noise, all the images before bleaching were averaged. The mean of the 1% of the total pixels with the lowest intensities was calculated as a normalization factor and all the images in the stack were then corrected by multiplication with the normalization factor and dividing by the mean image so that the overall intensities of all pixels in images were uniform. After background correction, the centers of the bleached regions and an appropriate unbleached reference were manually selected. The fluorescence recovery over time was fitted with the mathematical solution described in (18) in order to calculate the diffusion coefficients of the lipids in the SLBs. The mean of all individual recovery curves from different regions of interest was then fitted again to reduce the effect of random fluctuations. As a result, the mean diffusion coefficient for lipids in a SLB was obtained. The error was estimated by calculating the standard error of the mean of all individual fits.

For determining the diffusivity of membrane-anchored rKin430-SBP-GFP, single molecules were tracked using Fluorescence Image Evaluation Software for Tracking and Analysis (FIESTA) (21). Displacement data for all the single molecule trajectories were calculated for discrete time points and the displacement data were cumulated to calculate the average mean-square displacement (MSD) for every discrete time point. The first 8 points (based on the estimation published in (25)) of the MSD thus obtained were then fitted with a linear curve using the error bars as weights to get the diffusion coefficient of rKin430-SBP-GFP on SLBs.

For stepping motility assays, the mean velocity for individual motors was determined by calculating the slope of the trajectories in a kymograph (spacetime plot of intensity over a specified area, space dimension was chosen by a line drawn over a microtubule). Kymograph evaluation was performed using FIESTA software.

For membrane-anchored gliding motility assays, microtubules were tracked using FIESTA. All the connected tracks obtained from the software were visualized to manually exclude erroneous tracks from further analysis. Erroneous tracking could be due to, for example, microtubules crossing each other, sample drift or microtubule fragmentation. Therefore, the microtubule length was used as a control parameter for post-processing of tracks as the length is expected to remain constant over the duration of the experiment. The ensemble-average microtubule velocity for a particular surface motor density or diffusivity was obtained by calculating the MSD of the microtubule centers as a function of time. Due to the imaging of discrete frames, the time *t* is given as multiples *n* of the acquisition time interval such that *t* = *n*Δ*t*. The MSD is calculated for the non-overlapping time intervals using the formalism described in (25). The mean translational velocity and the diffusion coefficient for an individual microtubule were calculated by fitting the first 25 points of the MSD time plot with equation [1]. The MSD data for the fit was weighted by the inverse of the error. Only microtubules, which were in the field of view for more than 30 frames were analyzed. To calculate the mean velocity of an ensemble of microtubules at a particular motor density, we calculated the cumulated MSD for all the microtubules in an image stack. The cumulated MSD was calculated by cumulating the displacement data of all the individual microtubule for each discrete time point. The first 25 data points of the MSD data thus obtained were fitted with equation [1], to get an average microtubule gliding velocity. The MSD data for the fit was weighted by the inverse of error. The error for the fit was estimated by performing a boot strapping analysis (42). The MSD data for different microtubules were randomly picked (with replacement) keeping the total number of microtubule tracks analyzed the same as in the initial dataset. The standard deviation of the mean velocity obtained from the bootstrap analysis, gave the standard error of the mean of the initial dataset. Microtubule gliding velocities as function of motor density were fitted to equation [11] for different diffusivities of kinesin-1 on SLBs using the MATLAB curve fitting toolbox (nonlinear least square fitting with Lavenberg-Marquardt method). The fits were weighted by the inverse of the ensemble microtubule velocity error.

**Determination of the motor surface density on SLBs.** With the singlemolecule sensitivity of our TIRF set-up we were able to determine the actual number of diffusing motors directly by counting. As compared to the indirect measurement based on the total fluorescence intensity of GFP molecules, which can be skewed by several factors such as TIRF angle, optical aberrations, GFP clusters in the sample etc.; this direct measurement is more robust. Specifically, the kinesin-1 motor densities on biotinylated SLBs (at different bulk motor concentrations) were determined by incubating the SLBs with rKin430-SBP spiked with rKin430-SBP-GFP in a molar ratio 150:1. To avoid aberrations in counting due to photobleaching, the sample was focused by imaging the Atto 647n doped SLB before activating the perfect-focus mechanism of the Nikon TE 2000 Eclipse microscope. Movie streams with 50 frames at 100 ms exposure times were then recorded at five different fields of views, by exciting the sample with a 488 nm laser. The number of diffusing rKin430-SBP-GFP in the first three frames of the image stacks were then counted using the cell counter plug-in of the image processing and analysis software FiJi. The means and standard deviations of the measurements were then calculated. Gliding motility assays, to record the microtubule gliding velocity at that particular motor density, were then performed.

## Acknowledgements

We are grateful to C. Bräuer for technical assistance as well as C. Herold, L. Mu and M. Strempel for help with initial SLB experiments. We thank M. Braun and all other members of the Diez Lab for fruitful discussions. We acknowledge support from the Volkswagen Foundation (Grant I/84 087), the European Research Council (Starting Grant 242933 to SD), the Dresden International Graduate School for Biomedicine and Bioengineering (stipend to RG) as well as the German Research Foundation within the Cluster of Excellence Center for Advancing Electronics Dresden.

## Supporting Information

**Figure S1.**
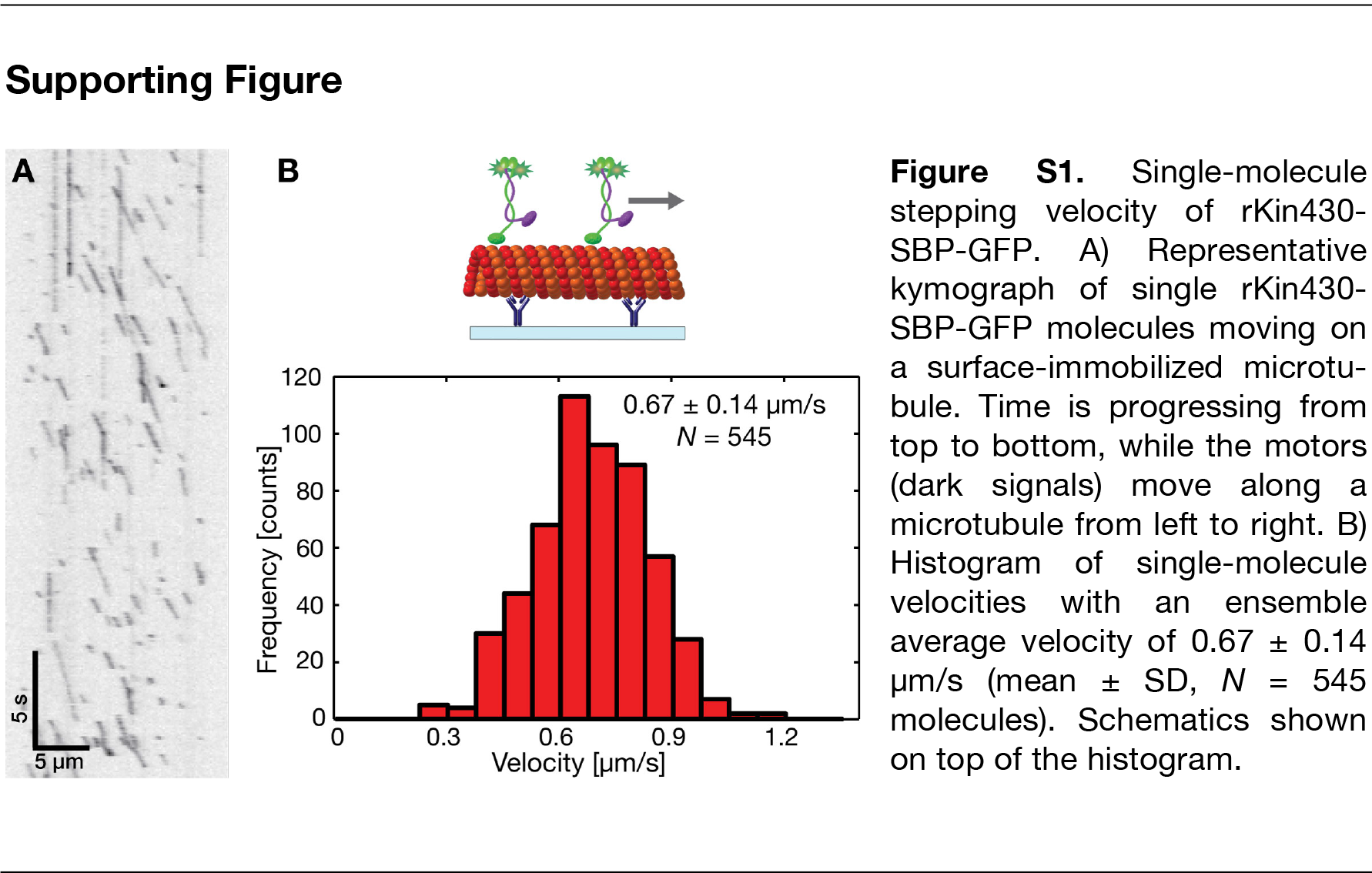
Single-molecule stepping velocity of rKin430-SBP-GFP. A) Representative kymograph of single rKin430-SBP-GFP molecules moving on a surface-immobilized microtubule. Time is progressing from top to bottom, while the motors (dark signals) move along a microtubule from left to right. B) Histogram of single-molecule velocities with an ensemble average velocity of 0.67 ± 0.14 μm/s (mean ± SD, *N* = 545 molecules). Schematics shown on top of the histogram.

### Supporting Movies

**Movie S1.** Membrane-anchored gliding motility at different motor densities with bulk rKin430-SBP concentrations 0.63 μM (top left panel, S1.1), 0.13 μM (top right panel, S1.2), 0.07 μM (bottom left panel, S1.3), and 0.13 μM in non-hydrolysable ATP analog AMP-PNP (bottom right panel, S1.4; Scale bar, 10 μm). Microtubule gliding velocity decreased with decreasing motor concentration and the respective trajectories became wigglier. In the presence of AMP-PNP, the microtubules anchored to SLBs via rigor motor-binding were diffusing only, without any linear translocation (S1.4).

**Movie S2.** rKin430-SBP gliding motility assay on different substrates: Motors anchored to SLB (left panel, S2.1) and motors immobilized on glass substrate (right panel, S2.2; Scale bar, 10 μm).

**Movie S3.** Addition of cholesterol to the DOPC SLBs reduced the diffusivity of lipids. Time-lapse movie showing FRAP for two different lipid compositions: DOPC:Cholesterol:DSPE-PEG-2000-Biotin = 79:20:1 (20% CH, left panel, S3.1) and DOPC:Cholesterol:DSPE-PEG-2000-Biotin = 39:60:1 (60% CH, right panel, S3.2). Fluorescence recovery of the photobleached region is faster in the SLB with 20% CH as compared to 60% CH (Scale bar, 10 μm).

**Movie S4.** Membrane-anchored gliding motility at different SLB diffusivities: 20 % CH (left panel, S4.1) and 60 % CH (right panel, S4.2). While microtubules do not cross each other when the membrane viscosity is low (left panel) they can cross each other at higher membrane viscosity (right panel, Scale bar, 10 μm).

